# Epigenetic Modifier Kdm6a/Utx Controls the Specification of Hypothalamic Neuronal Subtypes in a Sex-Dependent Manner

**DOI:** 10.1101/2022.07.25.501459

**Authors:** Lucas E. Cabrera Zapata, María Julia Cambiasso, Maria Angeles Arevalo

## Abstract

Kdm6a is an X chromosome-linked H3K27me2/3 demethylase that promotes chromatin accessibility and gene transcription and is critical for tissue/cell-specific differentiation. Previous results showed higher *Kdm6a* levels in XX than in XY hypothalamic neurons and a female-specific requirement for Kdm6a in mediating increased axogenesis before brain masculinization. Here we explored the sex-specific role of Kdm6a in the specification of neuronal subtypes in the developing hypothalamus. Hypothalamic neuronal cultures were established from sex-segregated E14 mouse embryos and transfected with siRNAs to knockdown Kdm6a expression (Kdm6a-KD). We evaluated by immunofluorescence the effect of Kdm6a-KD on Ngn3, a bHLH transcription factor regulating neuronal sub-specification in hypothalamus. Kdm6a-KD decreased Ngn3 expression in females but not in males, abolishing basal sex differences. Then, we analyzed Kdm6a-KD effect on *Ascl1, Pomc, Npy, Sf1, Gad1* and *Th* expression by RT-qPCR. While Kdm6a-KD downregulated *Ascl1* in both sexes equally, we found sex-specific results for *Pomc, Npy* and *Th. Pomc* and *Th* expressed higher in female than in male neurons and Kdm6a-KD reduced their levels only in females, while *Npy* expressed higher in male than in female neurons and Kdm6a-KD upregulated its expression only in females. Identical results were found by immunofluorescence for Pomc and Npy neuropeptides. Finally, using ChIP-qPCR we found higher H3K27me3 levels at *Ngn3, Pomc* and *Npy* promoters in male neurons, in line with Kdm6a higher expression and demethylase activity in females. These results indicate that Kdm6a plays a key sex-specific role in controlling the differentiation of neuronal populations regulating food intake and energy homeostasis.

## INTRODUCTION

Obesity and its associated comorbidities such as type 2 diabetes mellitus, cardiovascular disease, dyslipidemia and chronic inflammation, among others, have become a global health threat and a major socioeconomic burden for modern societies with a tendency towards sedentary lifestyles and diets based on hypercaloric/ultra-processed food products (Martin-Rodriguez et al., 2015; Khanna et al., 2022; Lustig et al., 2022). Although it is currently well known that significant sex differences exist in the prevalence, incidence and severity of a broad range of diseases, including obesity and obesity-related diabetes and cardiovascular disease (Seidell, 2005; Global et al., 2016; Hales et al., 2020; Teufel et al., 2021; Tsao et al., 2022), these differences are not well understood due to a combination of the lack of inclusion of sex as an influencing factor in preclinical studies and the underrepresentation of women in clinical trials. Consequently, there is an urgent international call for future research in the area and a better understanding of sex differences and their mechanistic underpinnings in health and disease (Vasquez-Avila et al., 2021; Regensteiner and Reusch, 2022).

The brain plays a pivotal role in the regulation of energy homeostasis. Particularly, the hypothalamus, one of the most sexually dimorphic brain structures (McEwen et al., 1979; McEwen, 1981; Bao and Swaab, 2010; Pfaff and Christen, 2013; Krause and Ingraham, 2017), is actively involved in the central control of feeding and energy expenditure, sensing and integrating peripheral metabolic and hormonal signals such as glucose, fatty acids, stomach-secreted ghrelin, pancreas-derived insulin and adipocyte-derived leptin to regulate food intake, glucose metabolism and energy balance, thereby ensuring that the nutrient demands of the body are fulfilled (Coll and Yeo, 2013; Timper and Bruning, 2017; Uwaifo, 2021). Two functionally antagonistic types of neurons in the arcuate nucleus of the hypothalamus have been identified as major coordinators of these processes: the neuropeptide Y (Npy) and agouti-related peptide (AgRP)-expressing neurons (Npy+ neurons) and the proopiomelanocortin (Pomc)-expressing neurons (Pomc+ neurons; Vohra et al., 2022). Npy+ neurons are orexigenic, meaning they stimulate appetite/hunger and promote feeding behavior (Leibowitz, 1991; Gropp et al., 2005), whereas Pomc+ neurons are anorexigenic, meaning they suppress appetite and promote satiety (Balthasar et al., 2005; Mountjoy, 2015). The appropriate function of these arcuate neuronal circuits and their coordination with other hypothalamic nuclei and extrahypothalamic brain regions also regulating metabolic homeostasis and feeding behavior critically depend on their correct shaping and integration during brain development, the disruption of which, at different levels, can cause a deficient system for the centra control of energy balance often leading to feeding-related diseases such as obesity, type 2 diabetes and metabolic syndrome, among others (Bingham et al., 2008; Gautron et al., 2015; Timper and Bruning, 2017; Farooqi, 2022). Hypothalamic development during embryogenesis is tightly controlled by basic Helix-Loop-Helix (bHLH) domain-containing transcription factors, which act in a temporally-coordinated sequential cascade to determine the neuronal fate and subsequently promote neuronal differentiation (Huang et al., 2014; Bedont et al., 2015; Baker and Brown, 2018). In this cascade of bHLH genes, proneural *Ascl1* (also known as *Mash1*) is expressed first and is required for the subsequent downstream expression of *neurogenin 3* (*Ngn3*; McNay et al., 2006; Pelling et al., 2011; Aujla et al., 2013) which, in turn, regulates the specification of different neuronal subtypes in the hypothalamus, including the Th+ (dopaminergic tyrosine hydroxylase-expressing neurons), Sf1+ (steroidogenic factor 1-expressing neurons), Pomc+ and Npy+ neuronal lineages (Pelling et al., 2011; Anthwal et al., 2013). Lysine demethylase 6A gene (*Kdm6a*, also known as *Utx*) is located on the X chromosome and encodes a histone demethylase involved in chromatin remodeling. By removing repressive di- and trimethyl (me2/3) groups on lysine (K) at position 27 in histone 3 (H3K27me2/3), Kdm6a promotes chromatin accessibility and allows gene expression (Hong et al., 2007; Tran et al., 2020). In XX individuals, it has been demonstrated in different cell types and developmental stages that *Kdm6a* consistently escapes X chromosome inactivation in both mice and humans and shows transcription from both alleles, leading to higher expression in females (Greenfield et al., 1998; Xu et al., 2008; Armoskus et al., 2014; Berletch et al., 2015; Tukiainen et al., 2017; Davis et al., 2020). Consistent with this, we have previously reported higher levels of *Kdm6a* expression in XX than in XY hypothalamic neurons and a female-specific requirement for the demethylase in mediating increased axogenesis before brain exposure to gonadal hormones (Cabrera Zapata et al., 2021). While Kdm6a is dispensable for the maintenance of embryonic stem cells, it plays a critical role in the determination of neural stem cells and the subsequent differentiation of these pluripotent cells into neurons and glia (Wang et al., 2012; Lei and Jiao, 2018; Yang et al., 2019; Shan et al., 2020; Subhramanyam et al., 2020), with deletion of *Kdm6a* leading to impaired dendritic arborization, synaptic formation, electrophysiological activity and cognition (Tang et al., 2017; Tang et al., 2020), and loss-of-function mutations in *KDM6A* causing cognitive deficits in humans (Miyake et al., 2013; Van Laarhoven et al., 2015; Bogershausen et al., 2016; Faundes et al., 2021). Here, we investigated the role of Kdm6a in the specification of neuronal subtypes in the developing hypothalamus and its differential requirement in males and females, focusing mainly on Pomc+ and Npy+ neuronal populations as essential elements in the central control of food intake and energy homeostasis.

## MATERIALS AND METHODS

### Animals

CD1 mice raised in our in-house colony at the Instituto Cajal (CSIC, Madrid, Spain) were used for this study. Animals received water and food *ad libitum* and were kept in controlled macroenvironmental conditions of temperature at 22 ± 2°C and a 12 h light/12 h dark periodic cycle. Procedures for care, welfare and proper use of all experimental animals followed the European Parliament and Council Directive (2010/63/EU) and the Spanish regulation (R.D. 53/2013 and Ley 6/2013, 11^th^ June) and were approved by our Institutional Animal Care and Use Committee (Comité de Ética de Experimentación Animal del Instituto Cajal) and by the Consejería del Medio Ambiente y Territorio (Comunidad de Madrid, PROEX 134/17).

### Hypothalamic neuronal cultures

CD1 mouse embryos at 14 days of gestation (E14, defining E0 as the day of the vaginal plug) were used to establish primary hypothalamic neuronal cultures. Donor embryos age was specifically selected with the purpose of avoiding exposure of neurons to the peak in gonadal testosterone secretion during *in utero* development, which occurs in male mice around E17 (O’Shaughnessy et al., 2006). Pregnant females were sacrificed by cervical dislocation under CO_2_ anesthesia, and embryos were dissected from the uterus. Neurons were cultured separately according to sex by observing the presence/absence of the spermatic artery in the developing gonads of embryos. The ventromedial hypothalamic region was dissected out and stripped off the meninges. Blocks of tissue were incubated for 15 min at 37 °C with 0.5% trypsin (Gibco, USA) and then washed three times with Ca^2+^/Mg^2+^-free Hank’s Buffered Salt Solution (Gibco). Finally, tissue was mechanically dissociated to single cells in 37 °C warm culture medium and cells were seeded. The medium was phenol red-free Neurobasal (Gibco) to avoid “estrogen-like effects” (Berthois et al., 1986) and was supplemented with B-27, 0.043% L-alanyl-L-glutamine (GlutaMAX-I) and 1% antibiotic-antimycotic containing 10.000 U/ml penicillin, 10.000 µg/ml streptomycin and 25 µg/ml amphotericin B (Gibco). Cells were plated on 6-well plates (Falcon, USA) at a density of 500-1000 cells/mm^2^ or on 10 mm glass coverslips (Assistent, Germany) at a density of 800 cells/mm^2^ for RT-qPCR/ChIP-qPCR or immunofluorescence, respectively. The surfaces of glass coverslips and plates were pre-coated with 1 μg/μl poly-L-lysine (Sigma-Aldrich, USA).

### Small interfering RNA (siRNA) transfection

Neurons were transfected by electroporation or lipofection using a mixture of 4 different siRNA sequences targeting *Kdm6a* transcripts at a final concentration of 40 nM total RNA (ON-TARGETplus Mouse Kdm6a Set of 4 siRNA, Dharmacon, UK); procedures and knockdown efficacy were previously described and demonstrated (Cabrera Zapata et al., 2021). A non-targeting siRNA sequence (ntRNA; Dharmacon) was used as control and co-transfection with pmaxGFP (Lonza, Switzerland) was performed in all cases for transfected neurons identification. For immunofluorescence, neurons were transfected by lipofection at 3 days *in vitro* (DIV) with target siRNA or ntRNA using Effectene Transfection Reagent (Qiagen, Germany) according to the manufacturer’s instructions, and after 18 h of knockdown were fixed and immunolabeled. For gene expression analysis, neurons were transfected by electroporation before seeding with target siRNA or ntRNA using a 4D-Nucleofector X Unit and the corresponding P3 Primary Cell nucleofection kit (Lonza) according to the manufacturer’s instructions, seeded and incubated for 3 DIV until processing for RNA isolation.

### RNA isolation and Reverse Transcription quantitative real-time PCR (RT-qPCR)

Total RNA was extracted using TRIzol reagent (Invitrogen, USA), purified and quantified by spectrophotometry on NanoDrop One (Thermo Fisher Scientific, USA) as previously described (Cisternas et al., 2015). 1 µg of RNA per sample was reverse transcribed to cDNA in a 20 µl reaction using M-MLV reverse transcriptase (Promega, USA) and random primers (Invitrogen), following manufacturer’s instructions. qPCR reactions were performed on a 7500 Real-Time PCR System (Applied Biosystems, USA) using SYBR Green Universal PCR Master Mix (Applied Biosystems). Primers (Table 1) were designed using the on-line Primer-Basic Local Alignment Search Tool (Primer-BLAST; National Institutes of Health, USA), selecting primer pairs spanning an exon-exon junction to restrict amplification specifically to mRNA. All primers were verified to amplify with a 95-100% efficiency by performing 4-point calibration curves. Relative quantification of mRNA expression was determined with the ΔΔCt method, using the BestKeeper index (Pfaffl et al., 2004) calculated for each sample from the Ct values of *Rn18s* (18S rRNA) and *Rpl13a* as control housekeeping genes. Control male samples were used as reference group.

**Table 1.**
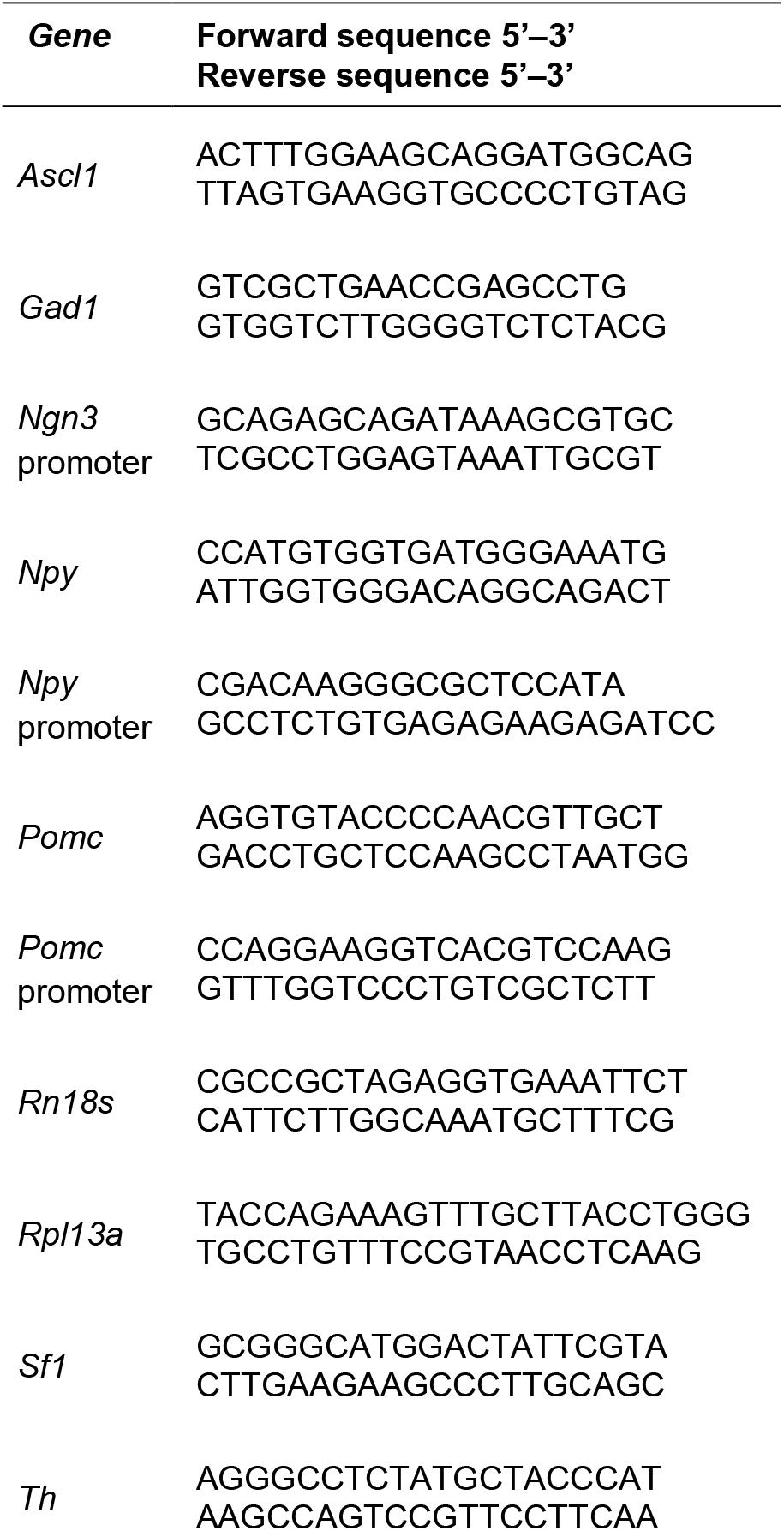
Primer sequences used for qPCR assays.

### Immunofluorescence

Neurons were fixed for 20 min at room temperature (RT) in 4% paraformaldehyde prewarmed to 37 °C, rinsed, permeabilized for 10 min with 0.5% Triton X-100 (Bio-Rad, USA) in phosphate buffered saline (PBS), blocked 1 h at RT in 1% BSA-PBS solution and incubated for 1 h at RT with the following antibodies diluted 1:1000 in 1% BSA-PBS: anti-Ngn3 mouse monoclonal antibody (F25A1B3, deposited by Madsen, O.D. to the Developmental Studies Hybridoma Bank, NICHD-NIH, maintained at The University of Iowa, Department of Biology, Iowa City, USA), anti-Pomc rabbit polyclonal antibody (H-029-30, Phoenix Pharmaceuticals, USA) or anti-Npy rabbit polyclonal antibody (T-4070, Peninsula Laboratories-BMA Biomedicals, Switzerland). After rinsing with PBS, cells were incubated for 1 h at RT with the secondary antibodies HRP goat anti-mouse (1:200 in 1% BSA-PBS) followed by 10 min amplification reaction with tyramide for the detection of Ngn3 (Tyramide Amplification Kid, 33003, Biotium, USA), or Alexa 594 donkey anti-rabbit for the detection of Pomc/Npy (1:1000 in 1% BSA-PBS; Jackson ImmunoResearch, USA). Finally, neurons were mounted on glass slides using gerbatol (0.3 g/ml glycerol, 0.13 g/ml Mowiol, 0.2 M Tris-HCl, pH 8.5) plus 1:5000 DAPI for nuclei staining.

### Imaging and quantitative image analysis

Imaging was done at 40x magnification using a standard Leica DMI 6000 fluorescence microscope (Leica, Germany) equipped with a digital camera of the same firm. To quantify fluorescent intensity, the soma of GFP-expressing neurons was outlined and the area and integrated density were measured in the corresponding channel for Ngn3, Pomc or Npy immunofluorescence signal using Fiji-ImageJ software (NIH, USA; freely available at https://imagej.nih.gov/ij/). Background fluorescence was also measured for each neuron/image analyzed. From these values, corrected total cell fluorescence (CTCF) intensity was calculated using the following equation (Martin Fitzpatrick, University of Birmingham, UK, available at https://theolb.readthedocs.io/en/latest/):

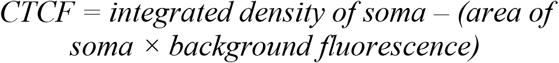

20-40 GFP-expressing neurons were randomly measured per experimental condition and culture (4 independent cultures).

### Chromatin immunoprecipitation (ChIP) analysis

ChIP assays were performed with ChIP-IT High Sensitivity Kit (Active Motif, USA). Briefly, ∼7-8 million hypothalamic neurons segregated by sex were cultured during 3 DIV and then fixed with fixation buffer containing 1.1% formaldehyde (Sigma-Aldrich). Cross-linked cells were scraped, centrifuged, washed 3 times with cold PBS and resuspended in ChIP buffer supplemented with PIC (Protease Inhibitor Cocktail; Active Motif) and PMSF (phenylmethylsulfonyl fluoride; Active Motif). Next, cells were homogenized using a dounce homogenizer with a tight-fitting pestle and the chromatin was sheared to ∼400-800 bp fragments by sonication with a Fisher Scientific Model 705 Sonic Dismembrator ultrasonic processor equipped with a microtip for small volume samples (FB705220, Thermo Fisher Scientific). 1% of total sonicated chromatin was kept as the input DNA and also used to determine DNA concentration at 260 nm on NanoDrop One (Thermo Fisher Scientific) for each sample. ChIP reactions were carried out overnight at 4 °C with 3 µg of sheared chromatin and 10 μg of anti-H3K27me3 rabbit polyclonal antibody (39155, Active Motif) or anti-mouse IgG as negative control (115-001-003, Jackson ImmunoResearch) in ChIP buffer supplemented with PIC. Immunoprecipitated DNA was quantified by real-time qPCR using primer pairs for the promoter regions of *Ngn3, Pomc* and *Npy* (Table 1) and the SYBR Green master mix and device detailed above. ChIP-qPCR data were normalized to the input DNA and expressed as percent of input.

### Statistical analysis

Data are presented as mean ± SEM and were statistically evaluated by two-way analysis of variance (ANOVA) with treatment/ChIP and gonadal sex as independent variables. Statistical significance of the effects of each independent variable and their interactions was tested. *Post hoc* comparisons of means by Fisher’s Least Significant Difference (LSD) test were performed for those variables/interactions for which ANOVA p-values were statistically significant. Statistical analysis was performed entirely with Statistica 8 software (StatSoft Inc., USA). p < 0.05 was considered statistically significant. Sample size (n) is indicated in the figure legends and was 3-6 independent cultures for RT-qPCR/ChIP-qPCR experiments or 40-100 transfected neurons from at least 4 independent cultures for immunofluorescence analysis. The number of independent cultures corresponds to the number of pregnant mothers from which embryos were obtained.

## RESULTS

### Kdm6a is female-specifically required for the expression of higher levels of Ngn3 in female than in male hypothalamic neurons

We have previously reported higher levels of expression of *Kdm6a* in hypothalamic neurons carrying two X chromosomes compared to those carrying one X and one Y chromosome regardless of gonadal sex, as well as a requirement of this higher *Kdm6a* expression and H3K27 demethylation activity in females for the increased expression of *Ngn3* in female compared to male hypothalamic neurons at a transcriptional (mRNA) level (Cabrera Zapata et al., 2021). To determine whether Kdm6a is also required for the sexually dimorphic expression of Ngn3 at the protein level, hypothalamic neuronal cultures derived from sex-segregated E14 mice were transfected with siRNAs targeting Kdm6a transcripts and the effect of Kdm6a knockdown on Ngn3 protein expression was analyzed by immunofluorescence. Remarkably, consistent with results evaluating mRNA levels, Kdm6a silencing led to a significant decrease in Ngn3 protein levels only in female-derived neurons without affecting male cultures, abolishing the sex differences for Ngn3 expression observed under control conditions (Figure 1; two-way ANOVA, sex-treatment interaction effect: F(1, 305) = 45.705, p < 0.0000001).

**Figure 1.**
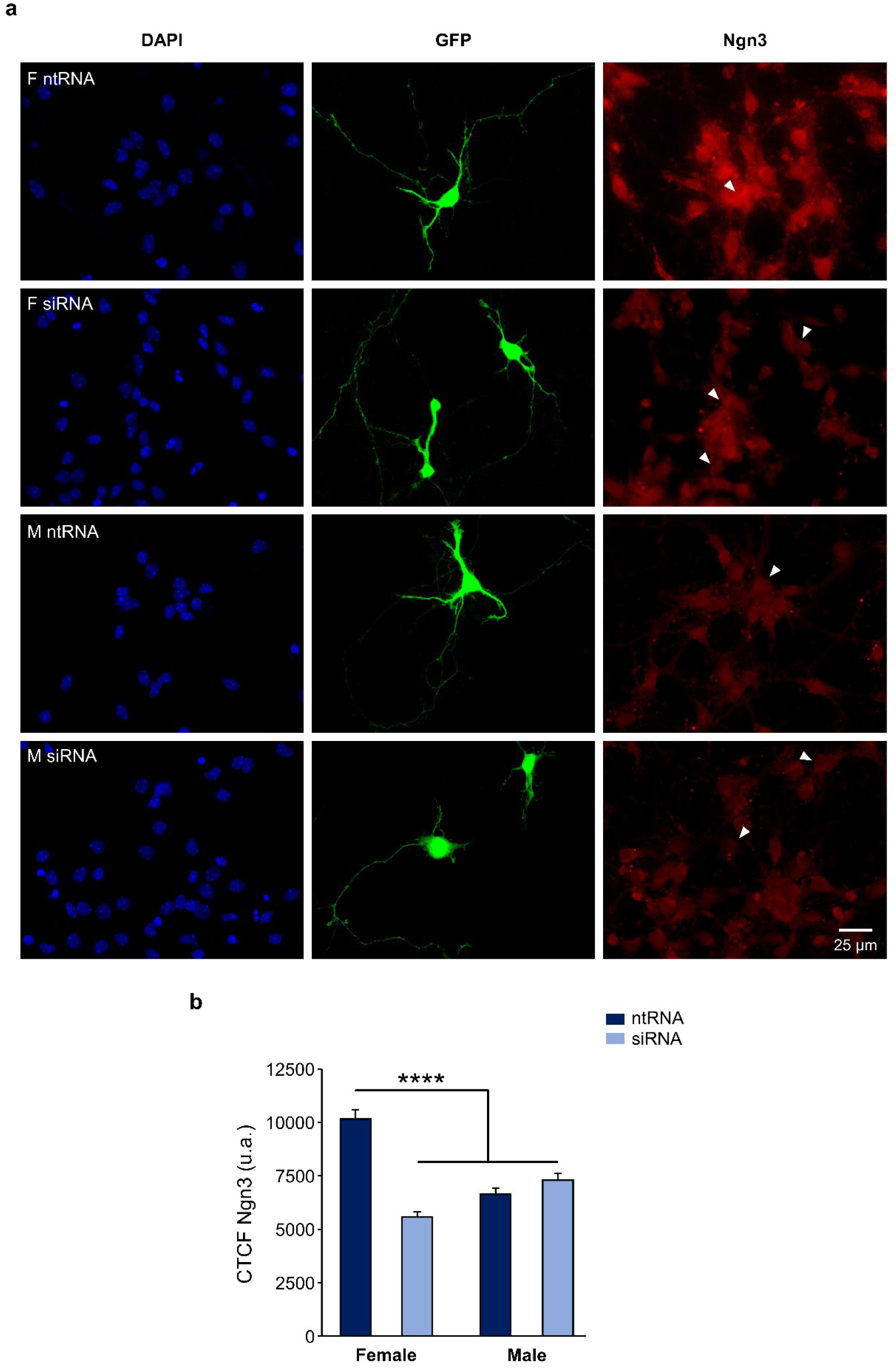
Kdm6a is required for the sexually dimorphic expression of proneural Ngn3. **(a)** Representative fluorescence images of female (F) and male (M) hypothalamic neurons co-transfected with a non-targeting siRNA sequence (ntRNA) and GFP or siRNA targeting Kdm6a (siRNA) and GFP at 3 DIV for 18 h. Ngn3 protein expression (red) was determined by immunofluorescence staining in transfected GFP+ neurons (green). Nuclei were stained with DAPI (blue). Arrowheads point to representative measured neuronal somas. **(b)** Quantification of fluorescence intensity for Ngn3 expressed as corrected total cell fluorescence (CTCF). Kdm6a knockdown by siRNA eliminated sex differences in Ngn3 protein expression by downregulating the proneural factor only in female-derived neurons. Data are mean ± SEM. n = 40-100 neurons from 4 independent cultures for each sex and treatment. ****p < 0.0001.

### Kdm6a is required for Ascl1 transcription and regulates the expression of Th, Pomc and Npy in a sexually dimorphic manner

Ngn3 has been shown to play an essential role in the specification of different neuronal subtypes in the hypothalamus, having opposing effects such as promotion of Pomc+ and Sf1+ and repression of Npy+ and Th+ fate (Pelling et al., 2011; Anthwal et al., 2013). However, such effects have not been properly addressed by considering sex as a crucial factor in hypothalamus development. Since we have demonstrated that Kdm6a has a sexually dimorphic expression pattern itself, and that it is specifically required in female-derived hypothalamic neurons to upregulate Ngn3 expression, then we analyzed by RT-qPCR the effect of Kdm6a downregulation by siRNAs on mRNA expression levels of *Ascl1, Pomc, Npy, Th, Gad1* and *Sf1*, all of them being molecular markers of different hypothalamic neuronal lineages (Romanov et al., 2020). While Kdm6a knockdown downregulated mRNA levels of the transcription factor *Ascl1* in both sexes equally (Figure 2; two-way ANOVA, treatment main effect: F(1, 15) = 10.05, p = 0.006), sex-specific effects were found for *Pomc, Npy* and *Th*. Interestingly, *Pomc* and *Th* were significantly more expressed in neurons derived from females than from males and Kdm6a silencing reduced their mRNA levels only in females (Figure 2; two-way ANOVA, sex-treatment interaction effect: Pomc: F(1, 14) = 5.76, p = 0.031; Th: F(1, 12) = 9.625, p = 0.009), while *Npy* expression was higher in male than in female-derived neurons in control conditions and Kdm6a knockdown upregulated its expression only in females (Figure 2; two-way ANOVA, sex-treatment interaction effect: F(1, 13) = 6.46, p = 0.025). No effects of sex or Kdm6a knockdown on *Sf1* or *Gad1* mRNA levels were found (Figure 2).

**Figure 2.**
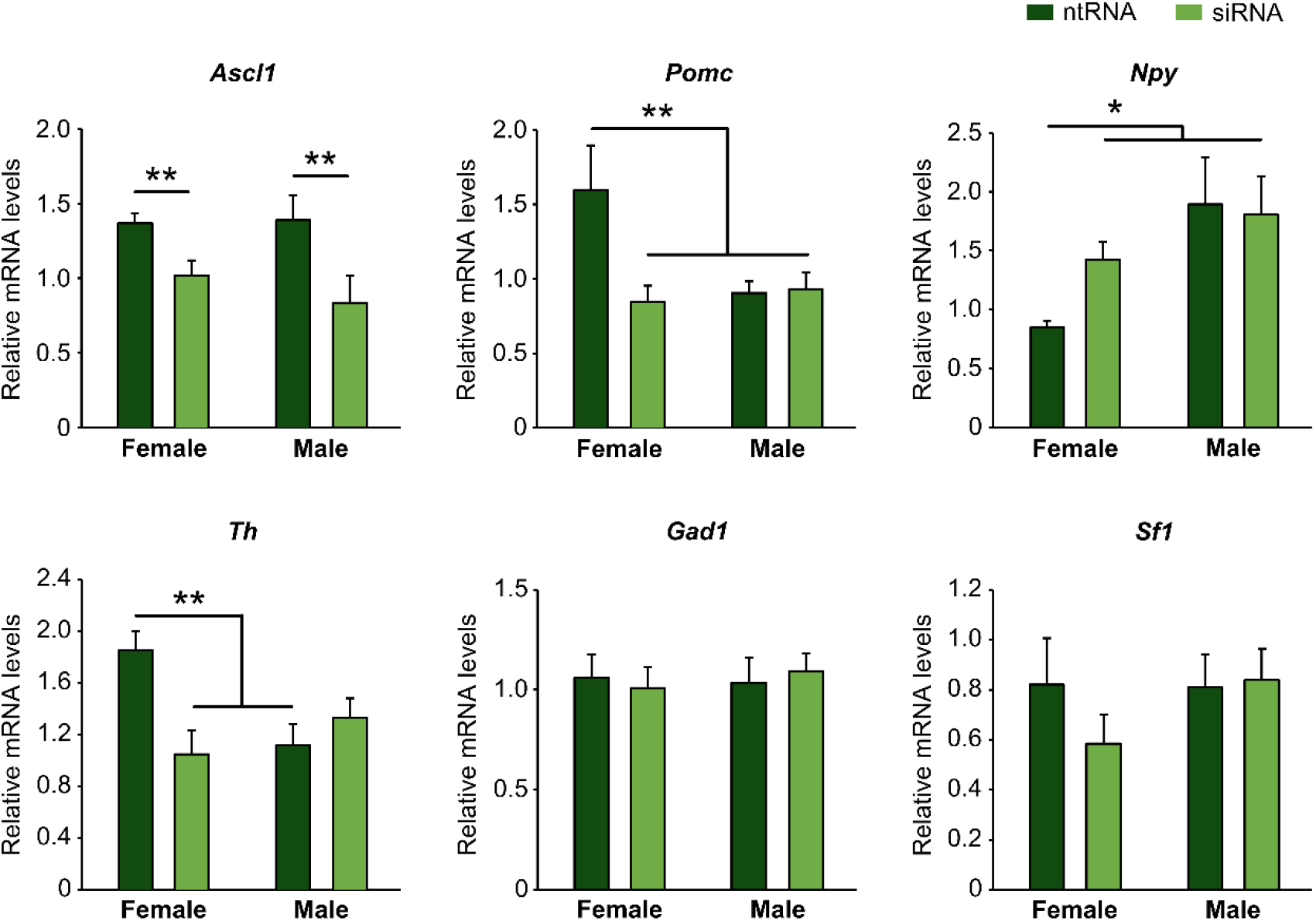
Kdm6a regulates gene expression of molecular markers of different hypothalamic neuronal lineages in a sex-specific manner. Kdm6a knockdown effect on *Ascl1, Pomc, Npy, Th, Gad1* and *Sf1* gene expression in male and female-derived hypothalamic neurons analyzed by RT-qPCR. Data are mean ± SEM. n = 4-6 independent cultures for each sex and treatment. *p < 0.05; **p < 0.01.

Having found sexually dimorphic mRNA expression patterns for *Pomc* and *Npy* regulated by Kdm6a in a sex-specific way, and considering the opposing roles of Pomc+ and Npy+ hypothalamic neurons regulating food intake and energy balance in mammals (Morton et al., 2006; Chen et al., 2022), we decided to analyzed the effect of Kdm6a silencing on Pomc and Npy protein expression levels by immunolabeling neurons co-transfected with siRNAs and pmaxGFP. Fluorescence intensity analysis for Pomc and Npy proteins in GFP-expressing hypothalamic neurons showed results consistent with the previous assessment at mRNA level. As shown in Figure 3, Pomc protein expression was significantly decreased after Kdm6a knockdown in female but not in male-derived hypothalamic neurons, erasing the sex differences in the expression levels of this neuropeptide observed in control conditions (two-way ANOVA, sex-treatment interaction effect: F(1, 263) = 10.64, p = 0.001). Regarding Npy, male hypothalamic neurons expressed significantly higher levels of this neuropeptide than female neurons under control conditions, whereas Kdm6a silencing had opposite effects in each sex, leading to increased Npy expression in females and decreased expression in males (Figure 4; two-way ANOVA, sex-treatment interaction effect: F(1, 282) = 9.44, p = 0.002).

**Figure 3.**
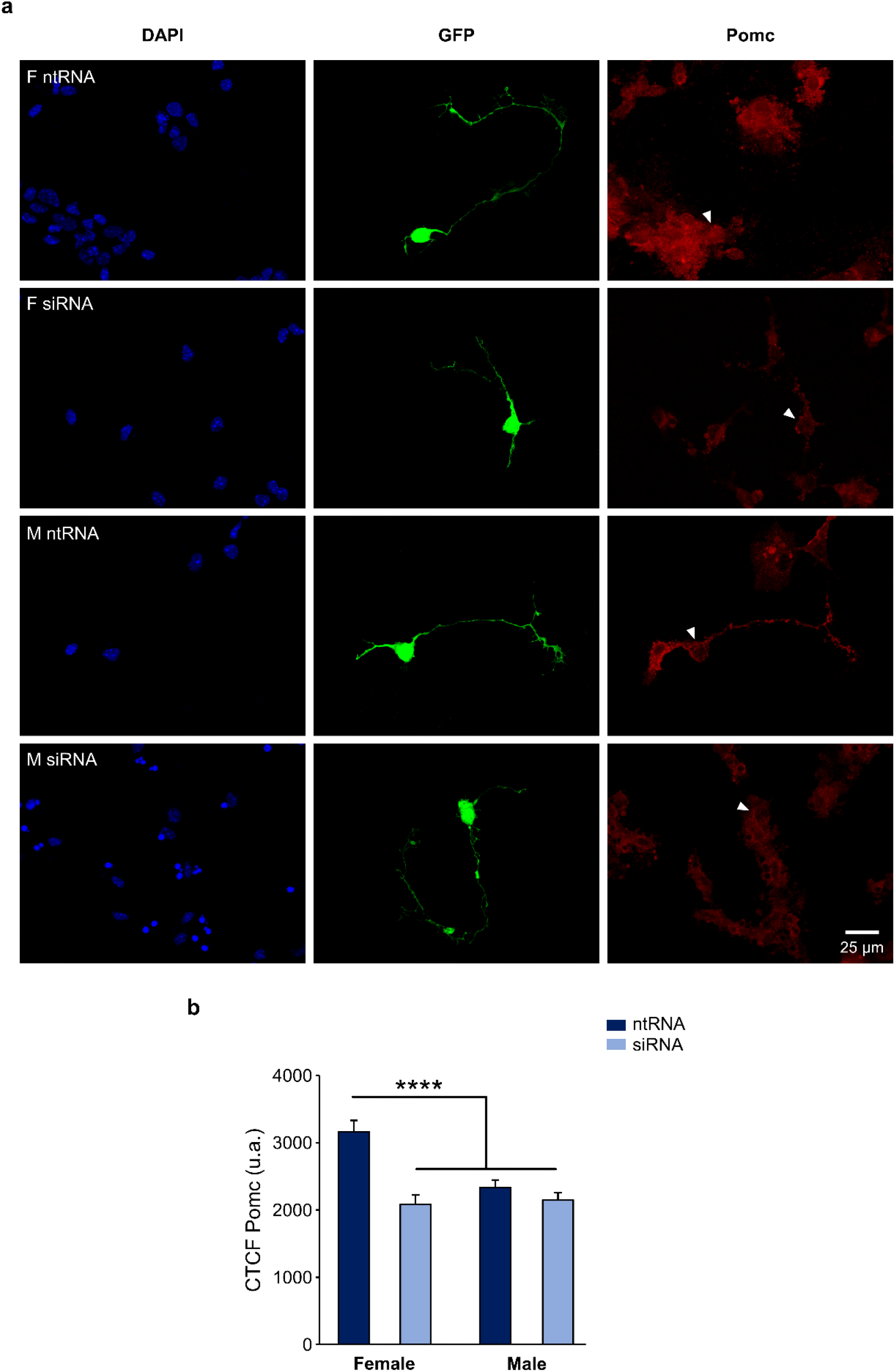
Kdm6a is required for Pomc sexually dimorphic expression. **(a)** Representative fluorescence images of female (F) and male (M) hypothalamic neurons co-transfected with a non-targeting siRNA sequence (ntRNA) and GFP or siRNA targeting Kdm6a (siRNA) and GFP at 3 DIV for 18 h. Pomc protein expression (red) was determined by immunofluorescence staining in transfected GFP+ neurons (green). Nuclei were stained with DAPI (blue). Arrowheads point to representative measured neuronal somas. **(b)** Quantification of fluorescence intensity for Pomc expressed as corrected total cell fluorescence (CTCF). Kdm6a knockdown by siRNA abolished sex differences in Pomc protein expression by downregulating the neuropeptide only in female-derived neurons. Data are mean ± SEM. n = 55-80 neurons from 4 independent cultures for each sex and treatment. ****p < 0.0001.

**Figure 4.**
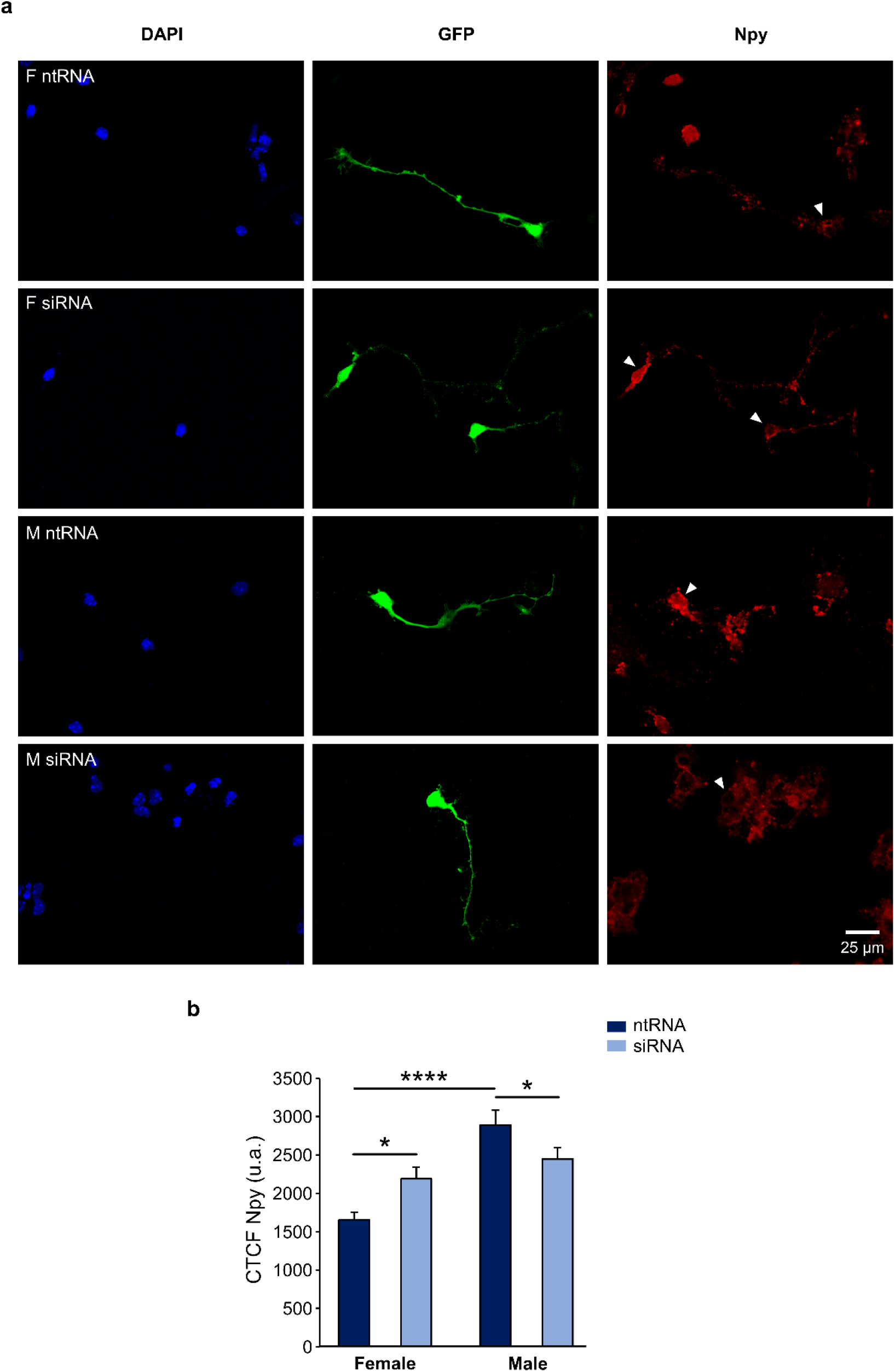
Kdm6a is required for Npy sexually dimorphic expression. **(a)** Representative fluorescence images of female (F) and male (M) hypothalamic neurons co-transfected with a non-targeting siRNA sequence (ntRNA) and GFP or siRNA targeting Kdm6a (siRNA) and GFP at 3 DIV for 18 h. Npy protein expression (red) was determined by immunofluorescence staining in transfected GFP+ neurons (green). Nuclei were stained with DAPI (blue). Arrowheads point to representative measured neuronal somas. **(b)** Quantification of fluorescence intensity for Npy expressed as corrected total cell fluorescence (CTCF). Npy expressed higher in male than in female hypothalamic neurons and Kdm6a knockdown by siRNA increased the neuropeptide expression in females while decreasing it in males. Data are mean ± SEM. N = 60-90 neurons from 4 independent cultures for each sex and treatment. *p < 0.05; ****p < 0.0001.

### *Ngn3, Pomc* and *Npy* promoters present higher binding levels to repressive H3K27me3 in male than in female hypothalamic neurons

Given the results clearly demonstrating that Kdm6a regulates Ngn3, Pomc and Npy expression differently in male and female hypothalamic neurons, and knowing that Kdm6a promotes gene transcription mainly by demethylating H3K27me3, we performed ChIP-qPCR assays in sex-segregated neuronal cultures to evaluate whether the promoter regions of *Ngn3, Pomc* and *Npy* genes are being controlled by the H3K27me3 histone epigenetic modification. Results expressed as qPCR data for promoter sequences in H3K27me3 or IgG (negative control) ChIP samples normalized to input (starting non-immunoprecipitated chromatin) confirmed the presence of the repressive H3K27me3 mark removed by Kdm6a at the promoters of *Ngn3, Pomc* and *Npy*, showing that the expression of these genes is regulated by H3K27 methylation/demethylation. Notably, in all cases we found significantly higher levels of promoter DNA bound to H3K27me3 in male than in female neurons, indicating a more suppressive effect for *Ngn3, Pomc* and *Npy* transcription in males (Figure 5; two-way ANOVA, sex-ChIP interaction effect: *Ngn3*: F(1, 9) = 6.29, p = 0.03; *Pomc*: F(1, 9) = 15.41, p = 0.003; *Npy*: F(1, 9) = 7.34, p = 0.02). These results are in line with the higher expression of Kdm6a and H3K27 demethylase activity in female than in male hypothalamic neurons previously reported (Cabrera Zapata et al., 2021).

**Figure 5.**
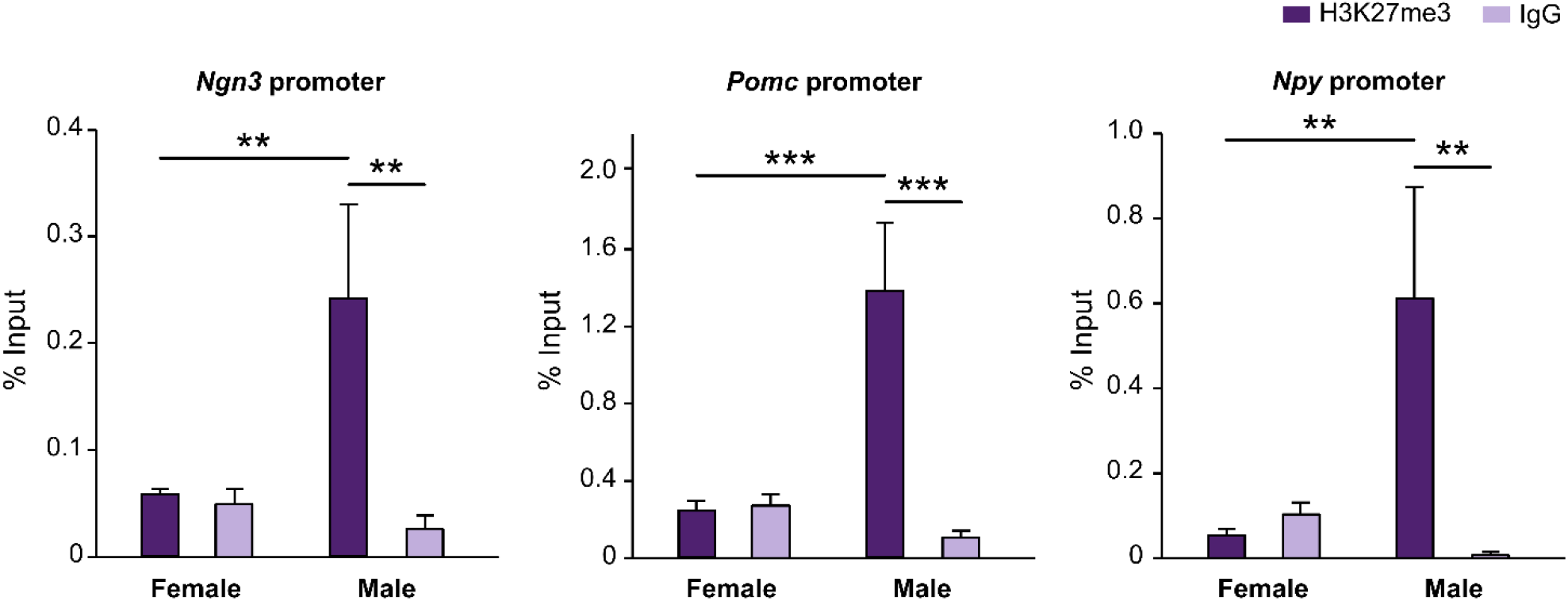
*Ngn3, Pomc* and *Npy* promoters are regulated by repressive H3K27me3 with a stronger suppressive effect in males. Levels of *Ngn3, Pomc* and *Npy* promoters DNA bound by H3K27me3 measured by ChIP-qPCR and expressed as % of input DNA. IgG antibody served as a negative control for H3K27me3 immunoprecipitation. Data are mean ± SEM. n = 3-4 independent cultures for each group. **p < 0.01; ***p < 0.001.

## DISCUSSION

In this work we present novel data suggesting a sex-specific role for the histone demethylase Kdm6a in the sexually dimorphic differentiation and specification of neuronal subtypes in hypothalamus. Kdm6a was female-specifically required for the expression of higher levels of the proneural transcription factor Ngn3 in female than in male hypothalamic neurons. Importantly, we have shown a Kdm6a sex-specific requirement for Pomc, Npy and Th sexually dimorphic expression in hypothalamic neurons. ChIP-qPCR data confirmed the presence of H3K27me3 repressive epigenetic marks regulating chromatin accessibility at *Ngn3, Pomc* and *Npy* promoter regions, indicating a role for Kdm6 demethylases in controlling transcription of these genes by removing H3K27 methylation. Notably, in all three genes, we found significantly higher levels of promoter DNA bound to H3K27me3 in male than in female neurons, indicating a more suppressive effect for their transcription in males. All these results clearly reveal dimorphic mechanisms for hypothalamic neuronal differentiation that depend on the X-linked epigenetic regulator Kdm6a and arise early in embryogenesis and before the critical action of gonadal hormones in the brain, thus contributing to the understanding of the role of sex chromosomes in brain sexual differentiation beyond the effects of gonadal hormones.

bHLH transcription factor Ngn3 plays an essential role in neuronal development and diversification in the hypothalamus. Performing genetic fate mapping studies and using *Ngn3*-null mutant mice (Gradwohl et al., 2000), it was possible to demonstrate that Ngn3+ progenitors contribute to subsets of arcuate Th+, Pomc+ and Npy+ neurons and ventromedial Sf1+ neurons, although the requirement for Ngn3 was different depending on the neuronal subtype: while Ngn3 promoted the development of Pomc+ and Sf1+ neurons, it inhibited the development of Npy+ and Th+ neurons (Pelling et al., 2011). Moreover, a conditional deletion of *Ngn3* specifically in the developing embryonic hypothalamus by using a *Nkx2*.*1iCre* transgenic mouse led in adulthood to a depletion of *Pomc* expression in arcuate neurons, a decrease in leptin sensitivity in ventral hypothalamic areas, and an associated obesity due to hyperphagia and reduced energy expenditure (Anthwal et al., 2013). However, although differences between male and female *Ngn3* conditional-knockout mice in parameters such as body weight gain and insulin and leptin insensitivity were reported (with more pronounced effects in females than in males; Anthwal et al., 2013), sex differences regarding Ngn3 role in the specification of hypothalamic neuronal subtypes have not been properly addressed. The hypothalamus is a brain region well known to be a highly sexually dimorphic structure (McEwen et al., 1979; McEwen, 1981; Pfaff and Christen, 2013), and our groups have already reported higher *Ngn3* expression levels in female than in male primary hypothalamic (Scerbo et al., 2014; Cisternas et al., 2020) and hippocampal (Ruiz-Palmero et al., 2016) neurons. In the present study, Kdm6a knockdown downregulated Ngn3 protein expression only in female hypothalamic neurons without affecting it in male neurons, clearly indicating a female-specific requirement of Kdm6a for higher Ngn3 levels in female than in male neurons. Furthermore, Kdm6a silencing affected the transcription of recognized molecular markers of different hypothalamic neuronal lineages, such as *Ascl1, Th, Pomc* and *Npy. Ascl1* (formerly known as *Mash1*) is another proneural bHLH transcription factor controlling both neurogenesis and neuronal subtype specification in hypothalamus (McNay et al., 2006; Alvarez-Bolado, 2019; Romanov et al., 2020), which has been reported to initiate a cascade of bHLH genes during early embryogenesis by inducing *Ngn3* expression that, in turn, promotes *Neurod1* and *Nhlh2* transcription downstream (McNay et al., 2006; Pelling et al., 2011). Our data show a requirement of Kdm6a for *Ascl1* mRNA expression equally important in male and female hypothalamic neurons, suggesting a key regulatory role for Kdm6a in neurogenesis and neuronal diversification in hypothalamus through the transcriptional control of a proneural master gene such as *Ascl1*. These results are not only relevant considering the proneural functions of Ascl1 during hypothalamic development *in utero*, but could also be of critical interest in the exciting, more recent and rapidly developing field of postnatal-adult hypothalamic neurogenesis (Kokoeva et al., 2005; Kokoeva 2007; Hourai and Miyata, 2013; Yoo and Blackshaw, 2018), where Ascl1 expression has been proposed to regulate the switch from the quiescent to the active state of neural stem cells (Sueda et al., 2019; Dou et al., 2021; Son et al., 2021). Additionally, we found higher mRNA levels of the dopaminergic molecular marker *Th* in female than in male-derived cultures and a female-specific requirement of Kdm6a for the expression of this sexually dimorphic pattern. Remarkably, it has recently been demonstrated that all Th+ dopaminergic neurons in hypothalamus are derived from Ascl1-expressing progenitors (Romanov et al., 2020) and that hypothalamic dopaminergic fate is modulated through the Ascl1/Ngn3 pathway (McNay et al., 2006; Pelling et al., 2011). Here we have shown a transcriptional regulatory role for Kdm6a at the three different levels of the Ascl1/Ngn3/Th axis, with sex-specific effects at Ngn3 and Th levels.

Results regarding the neuropeptides Pomc and Npy show sex-dependent expression patterns at both mRNA and protein levels, with Pomc expressing more in female than in male hypothalamic neurons and Npy showing the opposite pattern of higher expression in male than in female neurons. siRNA targeting Kdm6a mRNA decreased Pomc expression only in female neurons without affecting its levels in male neurons. On the other hand, Kdm6a downregulation increased Npy expression in female neurons while decreasing it in males. Although at first glance these results may seem controversial, they can be easily explained considering the higher Kdm6a and Ngn3 expression levels in female than in male neurons and the opposite roles of Ngn3 in promoting Pomc and repressing Npy fates previously reported (Pelling et al., 2011; Anthwal et al., 2013). Consequently, while in female neurons the higher levels of Kdm6a upregulate Ngn3 which, in turn, increases Pomc and decreases Npy expression, in male neurons the lower levels of Kdm6a and Ngn3 may lead to favor Npy over Pomc expression.

Ngn3 is well known as a master regulator of pancreatic and gut endocrine cell fate specification (Gradwohl et al., 2000; Jenny et al., 2002; Gasa et al., 2004; Mellitzer et al., 2010). While some studies have demonstrated a cooperation between Ngn3, H3K27 Kdm6 demethylases and other chromatin modifiers in the activation of downstream genes controlling pancreatic development (Fontcuberta-PiSunyer et al., 2018; Yu et al., 2018), no study to date has confirmed whether the promoter region of *Ngn3* is *per se* under regulation by H3K27 methylation marks, neither in pancreatic/enteroendocrine cells nor in neurons. As far as we are aware, our ChIP-qPCR data demonstrated for the first time that Ngn3, Pomc and Npy expression is regulated by the repressive H3K27me3 epigenetic motif controlled by Kdm6 histone demethylases. Noteworthy, we found significantly higher levels of H3K27me3 associated to the promoter regions of these genes in male than in female hypothalamic neurons, consistent with the higher expression and likely increased H3K27 demethylase activity of Kdm6a in females (Cabrera Zapata et al., 2021). These results together with results coming from mRNA and protein expression studies clearly show that Kdm6a controls the transcription of Ngn3, Pomc and Npy in hypothalamic neurons in a sex-dependent manner, while strongly suggesting this regulation occurs through H3K27 demethylation without discarding other mechanisms also displayed by Kdm6a such as promotion of H3K4 methylation and/or H3K27 acetylation by association to COMPASS/COMPASS-like complexes and histone acetyltransferases, respectively (Cho et al., 2007; Tie et al., 2012; Wang et al., 2012; Wang et al., 2017).

Taken together, results of the present study suggest a model whereby Kdm6a controls Pomc and Npy expression at the transcriptional and postranscriptional level in a sex-specific manner through H3K27 demethylation and regulation of the bHLH Ascl1/Ngn3 axis (Figure 6). On the one hand, female hypothalamic neurons, whose embryonic and perinatal development occurs in the absence of significant levels of gonadal hormones, show higher expression of the X-linked Kdm6a than male neurons (Cabrera Zapata et al., 2021). Higher Kdm6a levels in female neurons lead via H3K27 demethylation to an increase in Ngn3 levels which, in turn, could be mediating the higher Pomc and lower Npy levels observed in females compared to males through its opposite roles of promoting/repressing Pomc/Npy expression, respectively (Pelling et al., 2011). On the other hand, later in development during the so-called “critical period” of brain organization by gonadal hormones (E17-P10 in mice), developing testes start to secrete significant amounts of testosterone, leading to a perinatal exposure of male but not female neurons to the effects of 17β-estradiol (E2) converted intraneuronally from gonadal testosterone (Phoenix et al., 1959; Rhoda et al., 1984; O’Shaughnessy et al., 2006; McCarthy, 2008; Gagnidze et al., 2010). E2 organizational actions would upregulate Ngn3 expression in male hypothalamic neurons (Scerbo et al., 2014; Cisternas et al., 2020) which, subsequently, could lead to a potentiation of Ngn3-mediated effects promoting Pomc and repressing Npy expression. In addition, Kdm6a promotes Ngn3, Pomc and Npy transcription by removing repressive H3K27me3, with a stronger effect in females than in males consistent with the increased levels of the demethylase in females. Interestingly, while we have previously demonstrated that the expression pattern of Kdm6a with higher mRNA levels in females than in males does not change under different conditions of gonadal hormone secretion/exposure, such as different developmental stages in hypothalamus (before -E14-, during -P0- and after -P60-critical period) or under E2 treatment in hypothalamic neurons *in vitro* (Cabrera Zapata et al., 2021), a recent study has shown that Kdm6a colocalizes with estrogen receptor alpha (ERα) on a subset of ERα-target genes in an E2-dependent manner, cooperating with the receptor to upregulate the expression of these estrogen-responsive genes in a human breast cancer cell line (Xie et al., 2017). Further studies are needed to explore whether Kdm6a interacts with sex hormones and their receptors to mediate the sexually dimorphic neuronal differentiation and specification of neuronal subtypes in hypothalamus.

**Figure 6.**
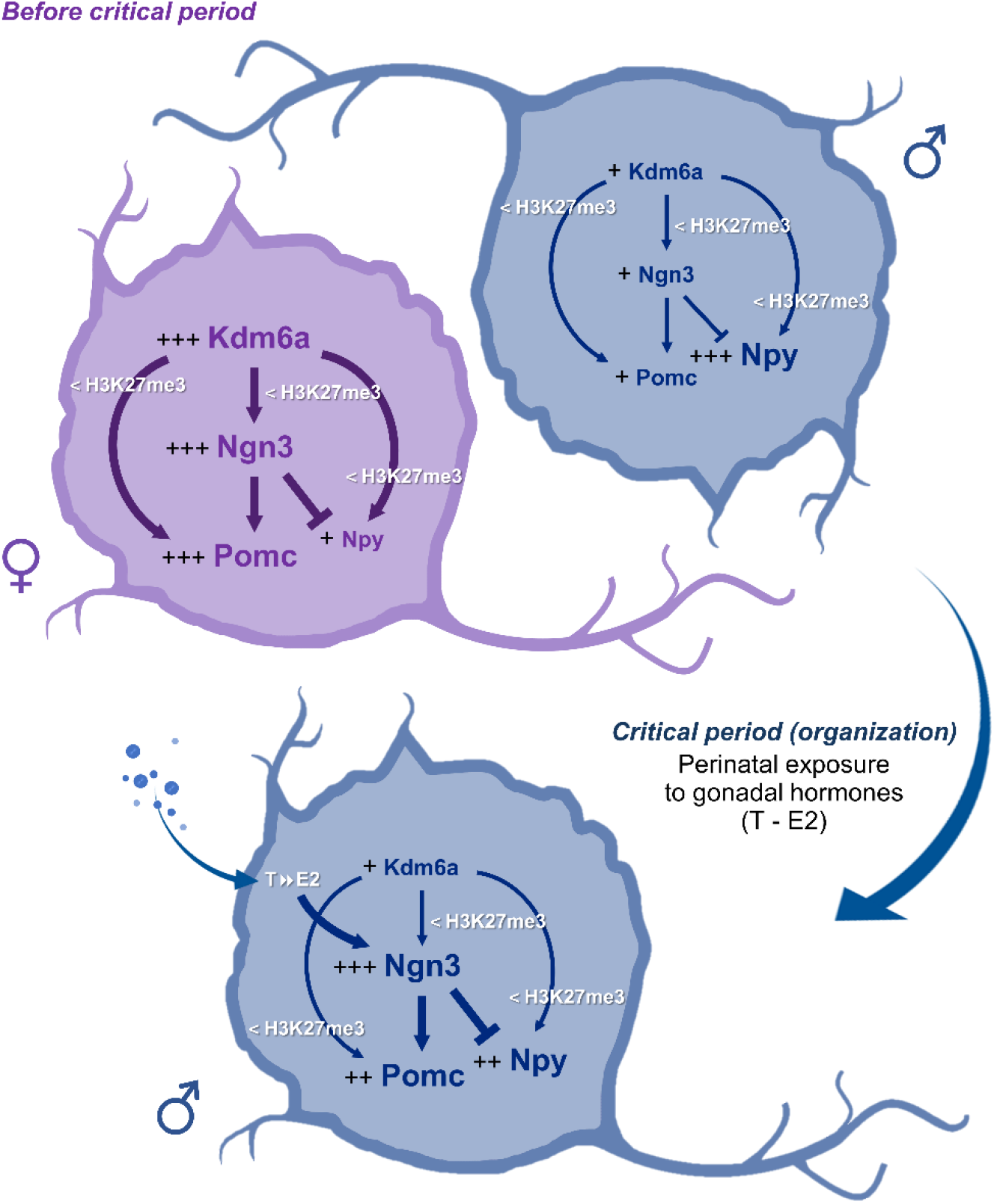
Hypothesis proposed to explain the sexually dimorphic mechanisms by which Kdm6a regulates the expression of Ngn3, Pomc and Npy in the developing hypothalamus. Before the critical period of hormonal organization of the brain, female hypothalamic neurons show higher expression of the X-linked Kdm6a than male neurons (Cabrera Zapata et al., 2021). Higher Kdm6a levels in female neurons upregulate Ngn3 transcription through H3K27me3 demethylation. In turn, Ngn3 is a proneural transcription factor that has been reported to promote Pomc and repress Npy expression (Pelling et al., 2011) and could be mediating the higher Pomc and lower Npy levels observed in females compared to males. In addition, Kdm6a promotes Ngn3, Pomc and Npy transcription by removing repressive H3K27me3, with a stronger effect in females than in males consistent with the increased levels of the demethylase in females. Later, during the critical period, male but not female neurons are exposed to the effects of estradiol (E2) converted intraneuronally from gonadal testosterone (T). E2 organizational actions increase Ngn3 expression in males (Scerbo et al., 2014; Cisternas et al., 2020), which could lead to increased Ngn3-mediated effects on Pomc and Npy expression. Comparisons between sexes are indicated as follows: more plus signs (+) and larger font size indicate higher expression levels; thicker arrows/lines indicate a stronger effect.

The current study highlights the importance of an X-linked gene with a sexually dimorphic pattern of expression, the genome-wide epigenetic modifier Kdm6a, in the development of the brain component of the feeding control circuit. We have demonstrated a critical requirement of Kdm6a for expression of the proneural bHLH Ascl1/Ngn3 transcriptional axis and the energy balance-related neuropeptides Pomc and Npy in a sex-dependent manner in hypothalamic neurons prior to the organizational actions of gonadal hormones on the developing brain, providing valuable new evidence for the impact of sex chromosomes on brain sexual differentiation beyond gonadal determination. Ngn3 is a developmental regulator of energy homeostasis systems in hypothalamus, pancreas and intestine, whereas Pomc+ and Npy+ hypothalamic neurons shape a central microcircuitry for the integration of peripheral signals providing feedback on energy status and for the control of feeding and energy expenditure, and disruptions or deficits in any of these systems have been associated with hyperphagia, obesity and obesity-related type 2 diabetes (Yaswen et al., 1999; Gradwohl et al., 2000; Anthwal et al., 2013; Bouret, 2017; Vohra et al., 2022). Notably, feeding and endocrine-related disorders such as feeding difficulties (some cases requiring long-term gastrostomy), gastrointestinal anomalies, tendency to overweight or obesity with age, hyperinsulinism, hypoglycemia and diabetes insipidus are commonly reported in Kabuki syndrome specifically caused by Kdm6a mutations, with male patients (carrying only one X chromosome and thus only one copy of Kdm6a) tending to be more severely affected (Lederer et al., 2014; Banka et al., 2015; Gole et al., 2016; Wang et al., 2019; Faundes et al., 2021). Delving deeper into the genetic, epigenetic and hormonal processes that regulate the development and function of the hypothalamus in each sex will lead to a better understanding of sexually dimorphic hypothalamic dysfunctions affecting energy homeostasis, allowing for more accurate treatments for epidemic diseases such as obesity in both men and women.

## STATEMENTS AND DECLARATIONS

### Author contributions

LECZ, MJC and MAA conceived and designed the research. LECZ performed all experiments, analyzed data, elaborated figures and wrote the first draft of the manuscript. All authors interpreted results of experiments. MJC and MAA edited and revised the manuscript. All authors read and approved the final manuscript. MAA drafted the manuscript.

## Acknowledgements

We thank the Secretaría General Iberoamericana (SEGIB), Fundación Carolina (Spain) and the International Brain Research Organization - Latin America Regional Committee (IBRO-LARC) for financially supporting part of Lucas E. Cabrera Zapata’s work at the Instituto Cajal, Madrid, Spain. We are grateful to Elisa Baides Rosell for her excellent technical assistance.

## Ethics approval

All experimental procedures with animals were approved and controlled by the Institutional Animal Care and Use Committee of the Cajal Institute (Comité de Ética de Experimentación Animal del Instituto Cajal) and by the Consejería del Medio Ambiente y Territorio (Comunidad de Madrid, PROEX 134/17), following the European Parliament and Council Directive (2010/63/EU) and the Spanish regulation (R.D. 53/2013 and Ley 6/2013, 11^th^ June).

